# Disruption of the Blood-Brain Barrier in 22q11.2 Deletion Syndrome

**DOI:** 10.1101/824987

**Authors:** Alexis M. Crockett, Sean K. Ryan, Adriana Hernandez Vasquez, Caroline Canning, Nickole Kanyuch, Hania Kebir, Guadalupe Ceja, James Gesualdi, Angela Viaene, Richa Kapoor, Naïl Benallegue, Stewart A. Anderson, Jorge I. Alvarez

## Abstract

Neuroimmune dysregulation is implicated in neuropsychiatric disorders including schizophrenia (SZ). As the blood brain barrier (BBB) is the immunological interface between the brain and the periphery, we investigated whether the BBB is intrinsically compromised in the most common genetic risk factor for SZ, the hemizygous deletion of chromosome 22q11.2 (22qDS). BBB-like endothelium (iBBB) differentiated from human 22qDS+SZ-induced pluripotent stem cells exhibited impaired barrier integrity, a phenotype substantiated in a mouse model of 22qDS. The proinflammatory intercellular adhesion molecule-1 (ICAM-1) was upregulated in 22qDS+SZ iBBB and 22qDS mice, indicating compromise of the BBB immune privilege. This immune imbalance resulted in increased migration/activation of leukocytes crossing the 22qDS+SZ iBBB. Finally, we found heightened astrocyte activation in murine and human 22qDS, suggesting that the BBB promotes astrocyte-mediated neuroinflammation. Overall, the barrier-promoting and immune privilege properties of the 22qDS BBB are compromised, and this might increase the risk for neuropsychiatric disease.

## INTRODUCTION

22q11.2 deletion syndrome (22qDS, also known as DiGeorge’s syndrome, velo-cardio-facial syndrome) is a genetic condition due to a 3 megabase hemizygous deletion of approximately 40 genes on chromosome 22 (Gur et al., 2017; Meechan et al., 2015). These patients present with cardiac malformations, facial deformities, gastrointestinal disturbances, and intellectual disability, as well as increased rates of neuropsychiatric conditions including attention deficits, autism spectrum disorder, and schizophrenia (SZ) (Gur et al., 2017). Remarkably, during adolescence and young adulthood roughly 25% of 22qDS patients develop SZ, a 25-fold increase in risk over the general population (Arinami, 2006; Karayiorgou & Gogos, 2004; Van, Boot, & Bassett, 2017). The 22q11.2 deletion includes junctional and mitochondrial genes that could influence barrier function (Arinami, 2006; Devaraju & Zakharenko, 2017; Greene et al., 2018; Greene, Hanley, & Campbell, 2019). However, the status of the blood brain barrier (BBB), the highly specialized central nervous system (CNS) endothelial cells, in 22qDS is unknown.

The vasculature of the BBB is composed of highly specialized endothelial cells characterized by specialized transport systems, higher mitochondrial volume fraction, and a paracellular cleft between adjacent endothelial membranes (Abbott, Rönnbäck, & Hansson, 2006; Obermeier, Daneman, & Ransohoff, 2013). The anatomical basis of barrier function is due to an elaborated network of junctional proteins composed of tight junctions (TJs) and adherens junctions (AJs) (Alvarez et al., 2015; Muldoon et al., 2013; Zihni et al., 2016). This network confers immune privilege to the CNS, by restricting interactions with the periphery (Abbott et al., 2010; Daneman & Prat, 2014; Engelhardt & Coisne, 2011; Muldoon et al., 2013). Additionally, immunoquiescent properties of the BBB endothelium also contribute to CNS immune privilege, specifically by downregulating expression of cell adhesion molecules, cytokines and chemokines to impede leukocyte transmigration into the CNS (Alvarez et al., 2015; Daneman & Prat, 2014). Interestingly, fluctuations of psychotic symptoms in SZ resemble the relapsing remitting pattern of prototypical immunological disorders, suggesting that the peripheral inflammation observed in SZ patients may contribute to disease progression (Eaton et al., 2006; Endres et al., 2015; Müller, 2018; Severance, Yolken, & Eaton, 2016). As the BBB stands as the interface between the CNS and the periphery (Abbott et al., 2010; Daneman & Prat, 2014), and because clinical data suggests BBB impairment in SZ (Aleksovska et al., 2014; Nishiura et al., 2017; Pollak et al., 2018), compromised BBB may contribute to disease progression (Greene et al., 2019; Kealy, Greene, & Campbell, 2018; Pollak et al., 2018). Therefore, we aimed to assess the barrier integrity and immunoquiescent properties of the CNS vasculature in the context of 22qDS.

Both barrier function and immune privilege of the BBB are modulated by the influence of astroglial processes ensheathing the CNS vasculature (Abbott et al., 2006; Alvarez, Dodelet-Devillers, et al., 2011; Cheslow & Alvarez, 2016). The close association of these perivascular astrocytes with the CNS vasculature positions them not only as critical cells to regulate CNS blood flow (Koehler, Roman, & Harder, 2009) and supply glucose to neurons (Verkhratsky & Nedergaard, 2018), but also the first cells to encounter breaches in the BBB (Alvarez, Cayrol, & Prat, 2011; Colombo & Farina, 2016). Astrocytes are key in the inflammatory cascade that results when the relative immune privilege status of the CNS is disrupted leading to neuroinflammation (Alvarez, Katayama, & Prat, 2013; Soung & Klein, 2018). As part of their role in the innate immune response, astrocytes become activated, producing cytokines and chemokines (Liddelow & Barres, 2017; Sofroniew, 2015). In light of the renewed emphasis on neuroinflammatory mechanisms in SZ (Miyaoka et al., 2017; Müller, 2018; Pollak et al., 2018; Severance et al., 2016; Wei & Hemmings, 2005) and as glial activation has been reported in the post mortem SZ brain (Catts et al., 2014; Colombo & Farina, 2016), we assessed neuroinflammation due to astrogliosis in 22qDS.

Here we have studied the barrier and immunological properties of the CNS vasculature in the context 22qDS. To this end, we employed an innovative approach based on the differentiation of human induced pluripotent stem cells (HiPSCs) from 5 different 22qDS patients with SZ (22qDS+SZ) and 5 age- and sex-matched healthy controls (HC) into BBB-like endothelium (iBBB). We found that the 22qDS+SZ iBBB exhibited substantially impaired barrier integrity, despite the genetic variability introduced by our approach. This phenotype was substantiated *in vivo* in a 22qDS mouse model harboring a homologous hemizygous deletion (Didriksen et al., 2017; Nilsson et al., 2018; Scarborough et al., 2019). We further interrogated junctional protein expression and immune privilege properties of the BBB *in vitro* and *in vivo*. We assessed the functional consequences of compromised BBB by determining the propensity of the 22qDS BBB to permit immune cell migration and activation *in vitro*, and promote perivascular astrocyte activation *in vivo*. Our results indicate that the 22q11.2 deletion reduces BBB integrity, alters the immune privilege of the CNS vasculature, and increases the transendothelial migration of peripheral immune cells, suggesting that BBB dysfunction may contribute to the increased susceptibility to SZ in 22qDS patients.

## MATERIALS AND METHODS

### Human induced pluripotent stem cells (HiPSCs)

HiPSC lines were generously received from Herbert M. Lachman, MD., Einstein University, Bronx, New York (15bc4, 553c2, 1804c6, 1bc4, 60c2, 30c1, and 3113c4) and Sergiu P. Paşca, MD., Stanford University, Stanford, California (1804.5, 2788.4, 511.1). All lines were transferred over to a feeder-free system with Stem MACS iPS-Brew XF media (Miltenyi Biotec 130-104-368). Lines were tested for mycoplasma using Lookout mycoplasma PCR detection kit (Sigma MP0035).

### Differentiation of iPSCs into blood-brain barrier endothelium

HiPSCs were differentiated into BBB-like endothelium following the protocol previously published (Hollmann et al., 2017; Lippmann et al., 2012). In brief, HiPSCs were plated onto Matrigel (Corning 354230) coated 6 well TC-treated plated (Falcon 353046) at 100,000 cells/well in Stem MACs iPS-Brew XF media. The following day (Day 0) media was fully exchanged (2 mLs) to unconditioned medium (UM): DMEM/Ham’s F12 containing 20% Knockout Serum Replacer (Invitrogen), 1× MEM nonessential amino acids (Invitrogen), 1 mM l-glutamine (Sigma), 0.1 mM β-mercaptoethanol (Sigma). Full exchanges were performed every day through Day 5. On day 6, media was fully exchanged (4 mLs) to endothelial cell medium (ECM): human Endothelial Serum-Free Medium (Invitrogen 11111) with 1% platelet-poor plasma-derived bovine serum (Biomedical Technologies 50-443-029) with 20 ng/mL bFGF (Peprotech 100-18B), and 10µM retinoic acid (RA; Sigma R2625). An addition 2 mLs of ECM with RA and bFGF was added on day 7. On Day 8, iBBB endothelial cells were frozen down. Previous publications have extensively validated the iBBB cells produced by this differentiation protocol (Hollmann et al., 2017; Lippmann et al., 2012); to determine that the 22q11.2 deletion did not affect the differentiation process, we confirmed similar expression levels in vasculature-specific molecules by flow cytometry (Supplementary Figure 2), and in junctional proteins by immunofluorescence (Supplementary Figure 1A) and western blot (Figure 2A).

### Cryopreservation of iBBB

On Day 8 of differentiation, iBBBs were washed twice with PBS (Phosphate-buffered saline) (1X), then lifted with StemPro accutase (ThermoFisher Scientific A11105-01) and spun down at 1,000 rpm for 5 min. The supernatant was aspirated, and the cells were re-suspended in 60% EC medium with retinoic acid, 30% FBS (fetal bovine serum) (Hyclone SH30071.03HI), and 10% DMSO at 2 x 10^6^ cells/mL. Cells were stored in liquid nitrogen.

### iBBB culturing

Plates were coated with a collagen/fibronectin mixture composed of 50% sterile H_2_O, 40% collagen from human placenta (Sigma C5533), and 10% fibronectin from bovine plasma (Sigma F1141) as previously described (Hollmann et al., 2017; Lippmann et al., 2012). Cells were counted and plated according to Table 1 in ECM (human endothelial Serum Free Media (Invitrogen) with 1% platelet-poor plasma-derived bovine serum (Alfa Aesar J64483)) with 20 ng/mL bFGF (Peprotech), 10 μM RA (Sigma), and 1:1000 Y27632 (ROCK inhibitor) (R&D 1254). Media was changed 24 hours later to ECM, and subsequently changed every 48 hours after, allowing cells to grow at 37°C in 5% CO_2_ until confluency was reached, for a maximum of 5 days. All experiments were conducted in paired format, in which 22qDS+SZ and age- and sex-matched HC iBBB endothelial cells were thawed, plated and analyzed in parallel. All iBBB data is presented as color/shape-matched pairs as follows: 15bc4/553c2, red circle; 30c1/3113c4, gray square; 60c2/511.1, blue diamond; 1804.5/2788.4, green triangle; 1804c6/1bc4, purple hexagon.

**Table 1.** iBBB culturing specifications.

| Experiment | Cell Density | Media Volume | Plating Substrate |
| --- | --- | --- | --- |
| TEER | $1.75 \times 10^5$ | 400 μL | 8W10E+ electrode arrays<br>(Applied Biophysics, Troy, NY, USA) |
| Immunofluorescence | $8 \times 10^4$ | 300 μL | 8 well chamber slide (ibidi) |
| Flow cytometry<br>Western blot | $1.8 \times 10^5$ | 2 mL | Nunclon™ Delta Multidishes, 6 well (ThermoFisher Scientific) |
| Monocyte migration | $8 \times 10^4$ | (Bottom) 1.5 mL, (Top) 600 μL | Falcon cell culture inserts<br>(ThermoFisher Scientific) |

### iBBB TEER

The electrical properties of confluent monolayers of iBBB endothelial cells were measured as previously described (Alvarez, Dodelet-Devillers, et al., 2011). In brief, the Electric Cell-substrate Impedance Sensing (ECIS) methodology was employed making using of the ECIS Zϴ instrument and 8W10E+ electrode arrays (Applied Biophysics, Troy, NY, USA). The TEER properties of iBBB cell monolayers were recorded at 2000 Hz over a period of 72 hrs. Resistance of the iBBB cell monolayers were assessed at confluency (determined as maximum resistance of the monolayer), and data is presented as fold change of the 22qDS+SZ iBBB, normalized to HC pair.

### iBBB immunofluorescence

Cells were washed twice with TBS (1X) (Tris-buffered saline) and fixed with 70% Ethanol for 5 min. Unspecific antibody binding was blocked for 30 min with 10% normal donkey serum (Sigma) followed by overnight 4°C incubation with rabbit polyclonal antibodies against claudin-5 (1:100; Life Technologies) and ZO-1 (1:200; Life Technologies) diluted in 3% normal donkey serum. After multiple washes in TBS (1X) with 0.025% Tween 20 (Amresco), secondary antibody Alexa Fluor® 594 AffiniPure Fab Fragment Donkey Anti-Rabbit IgG (1:300; Jackson Immunoresearch) was added and incubated for 60 minutes at room temperature. iBBB monolayers were then washed twice in TBS (1X) with 0.025% Tween 20 and mounted with Gelvatol containing Hoescht (1:1000; Invitrogen) for nuclear staining. Cells were imaged using a Leica SP5 confocal microscope (Leica Microsystems). Images were obtained using an Olympus IX83 set up for brightfield and fluorescence, and equipped with a motorized X, Y, Z stage and a spinning disk confocal head (X-Light V2, Crestoptics s.r.l., Rome, Italy) using a Hamamatsu R2 cooled CMOS camera (Hamamatsu City, Japan) operated by the MetaMorph software (Molecular Devices, LLC, Sunnyvale, CA).

### iBBB flow cytometry

Following 5 days in vitro, iBBB cells were detached with StemPro accutase (ThermoFisher Scientific A11105-01) and nonspecific binding was blocked with Mouse IgG Isotype Control (Invitrogen) for 25 minutes at 4°C. Cells were stained with monoclonal mouse anti-human antibodies as shown in Table 2 for 30 minutes at 4°C. Data was collected on LSRFortessa (BD Biosciences). Mean fluorescent intensity (MFI) of the ICAM-1^+^ population was measured with FlowJo version 10 software (BD biosciences). ICAM-1 expression in each donor pair was analyzed in 2-4 separate experiments.

**Table 2.**
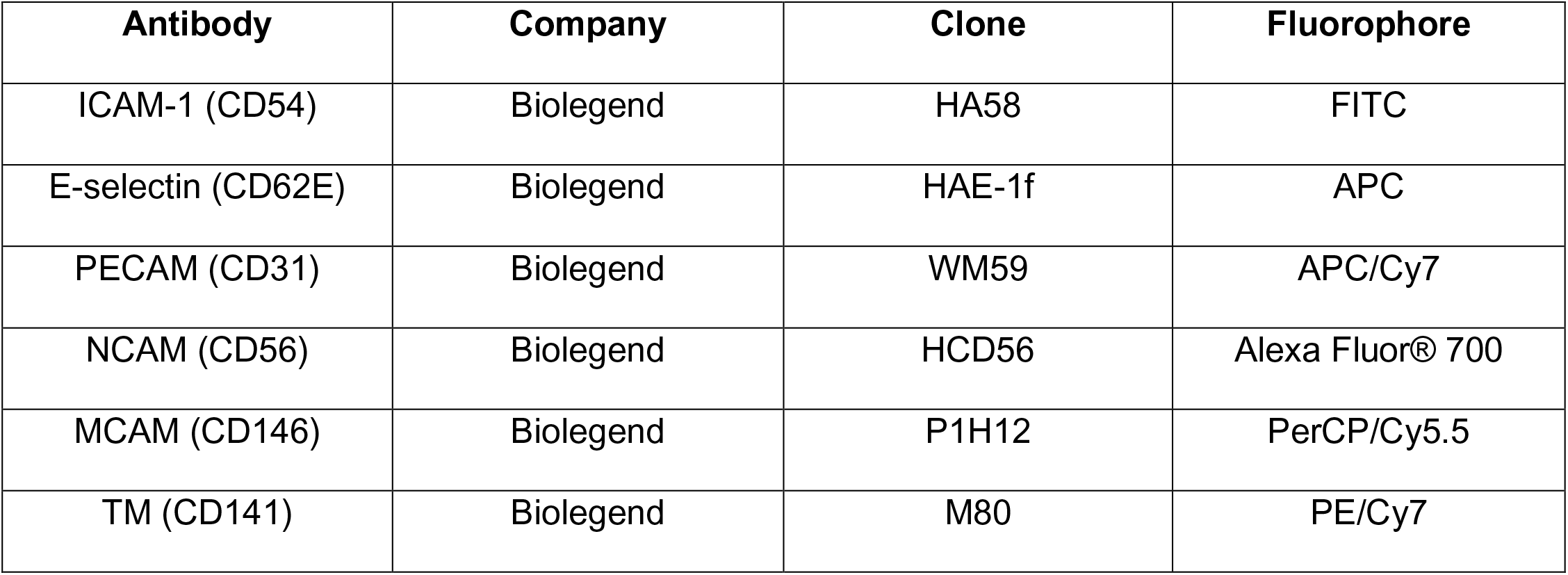
iBBB surface molecule antibodies.

| **Antibody** | **Company** | **Clone** | **Fluorophore** |
| --- | --- | --- | --- |
| ICAM-1 (CD54) | Biolegend | HA58 | FITC |
| E-selectin (CD62E) | Biolegend | HAE-1f | APC |
| PECAM (CD31) | Biolegend | WM59 | APC/Cy7 |
| NCAM (CD56) | Biolegend | HCD56 | Alexa Fluor® 700 |
| MCAM (CD146) | Biolegend | P1H12 | PerCP/Cy5.5 |
| TM (CD141) | Biolegend | M80 | PE/Cy7 |

### iBBB western blot

Following 5 days in vitro, cells were detached with StemPro accutase (ThermoFisher Scientific A11105-01), incubated in RIPA Lysis buffer (Amresco) with 1:100 protease inhibitor (Sigma) on ice for 30 minutes and subsequently centrifuged. Protein content was quantified using the Pierce™ BCA Protein Assay kit according to manufacturer protocols (Thermo Fisher Scientific). 20 μg of protein lysate was run on 4-20% Mini-PROTEAN TGX Precast Gel (BioRad), transferred to nitrocellulose membrane (BioRad) and blocked with Odyssey Blocking Buffer (Licor) for 1 hour at room temperature. Primary polyclonal rabbit antibodies against claudin-5 (1:500; Life Technologies) and ZO-1 (1:250; Life Technologies) were diluted in Odyssey Blocking Buffer and incubated overnight at 4°C. The membrane was washed with TBS (1X) with 1% Tween 20 (Amresco), secondary antibodies IRDye 680RD donkey anti-mouse and IRDye 800CW donkey anti-rabbit (Licor) were diluted 1:5000 in Odyssey Blocking Buffer, and incubated 1 hour at room temperature. Standard monoclonal mouse anti-β actin (1:10000; Sigma AC-74) was incubated 1 hour at room temperature. The membrane was developed on Odyssey Infrared Imaging System 9120 (Licor), and average pixel intensity of each band was measured by ImageJ.

### Immune cell transmigration across iBBB monolayers

To investigate monocyte migration across the 22qDS BBB, we utilized a transwell model in which iBBB cells were grown on the upper chamber of 6.4 mm filter Falcon cell culture inserts (ThermoFisher Scientific) for 5 days. Monocytes were isolated from the blood of 8 healthy volunteers as previously described (Kebir et al., 2009). In brief, peripheral blood mononuclear cells were isolated using density gradient centrifugation on Ficoll-Paque™ (GE Healthcare). CD14^+^ cells were obtained using the MACS isolation columns according to the manufacturer’s protocol (Miltenyi). CD14^+^ cells (1×10^6^ per well) were allowed to migrate for 18 hours, and the activation state of migrated monocytes was assessed using flow cytometry immunostaining. Nonspecific binding was blocked with Mouse IgG Isotype Control (Invitrogen) for 25 minutes at 4°C. Cells were stained with mouse anti-human antibodies shown in Table 3, diluted 1:100 for 30 minutes at 4°C. Data was collected on LSR Fortessa (BD Biosciences), and analysis was performed with FlowJo version 10 software (BD biosciences). Each HC/22qDS+SZ iBBB pair was run in triplicate in 2 different experiments, each time with monocytes isolated from a different healthy volunteer. Surface marker expression was evaluated within the CD14^+^ MHC-II^+^ positive population.

**Table 3.**
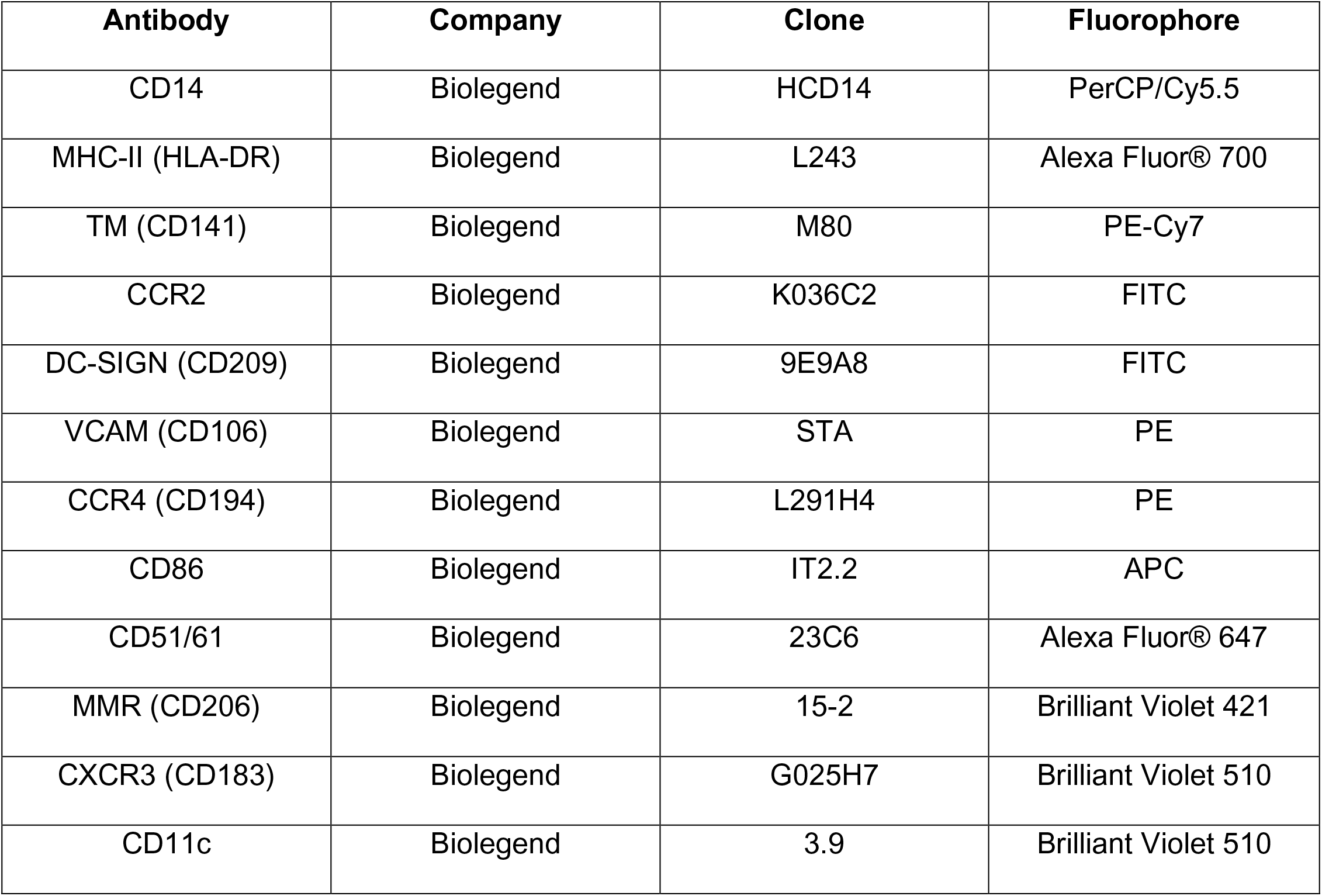
Monocyte migration antibodies.

| **Antibody** | **Company** | **Clone** | **Fluorophore** |
| --- | --- | --- | --- |
| CD14 | Biolegend | HCD14 | PerCP/Cy5.5 |
| MHC-II (HLA-DR) | Biolegend | L243 | Alexa Fluor® 700 |
| TM (CD141) | Biolegend | M80 | PE-Cy7 |
| CCR2 | Biolegend | K036C2 | FITC |
| DC-SIGN (CD209) | Biolegend | 9E9A8 | FITC |
| VCAM (CD106) | Biolegend | STA | PE |
| CCR4 (CD194) | Biolegend | L291H4 | PE |
| CD86 | Biolegend | IT2.2 | APC |
| CD51/61 | Biolegend | 23C6 | Alexa Fluor® 647 |
| MMR (CD206) | Biolegend | 15-2 | Brilliant Violet 421 |
| CXCR3 (CD183) | Biolegend | G025H7 | Brilliant Violet 510 |
| CD11c | Biolegend | 3.9 | Brilliant Violet 510 |

### Mice

All experiments were conducted in accordance with IACUC at the University of Pennsylvania. 4-6-month-old male and female 22qDS mice and WT littermates (Taconic) were anesthetized by either isofluorane or CO_2_ and transcardially perfused with ice cold PBS (1X). Brains were either frozen in Optimal Cutting Temperature (OCT) Compound (Fisher HealthCare) for sectioning, kept fresh in RPMI (1X)+GlutaMAX (Gibco) for flow cytometry or frozen at −80°C for western blotting.

Mouse anterior cortex was thawed and sonicated in RIPA Lysis buffer (Amresco) on ice using an Ultrasonic Homogenizer (Biologics Inc.). Western blot analyses proceeded as described above. N=11 WT, 10 22qDS (15 males, 6 females).

### Mouse CNS endothelial cell flow cytometry

Microvascular endothelial cells were isolated from the anterior portion of the CNS for flow cytometry analysis as previously described (Crouch & Doetsch, 2018). In brief, tissue was minced in HBSS (1X) (Hank’s Balanced salt solution, ThermoFisher Scientific) containing 1% bovine serum albumin (Fisher BioReagents), 0.1% glucose (Sigma) and 0.5 mg/mL DNase I (Roche) and was subsequently digested with 3 mg/mL Collagenase/Dispase (Roche) in PBS (1X) with 2% FBS (Serum Solutions International) for 30 minutes at 37°C with agitation. Following centrifugation, cells were resuspended in Trituration solution (PBS (1X) with 2% FBS and 0.5 mg/mL DNase I) and endothelial cells were obtained using density gradient centrifugation with 22% Percoll (GE Healthcare) in PBS (1X). Nonspecific binding was blocked with rat anti-mouse CD16/32 (1:100; Biolegend 93) for 20 minutes at 4°C, and cells were stained with rat anti-mouse ICAM-1 (1:100; Biolegend YN1/1.7.4 in PerCP), rat anti-mouse CD45 (1:100; BD Horizon™ 30-F11 in Brilliant Violet 510), rat anti-mouse CD102 (1:100; Biolegend 3c4 (MIC2/4) in Alexa Fluor® 488) and rat anti-mouse CD31 (1:100; Biolegend 390 in APC/Fire™ 750) for 30 minutes at 4°C. Cells were fixed in 4% PFA (paraformaldehyde) for 10 minutes at room temperature, and data was collected on LSRFortessa (BD Biosciences). MFI of the ICAM-1-positive endothelial (CD45^-^ CD102^+^ CD31^+^) population was measured with Flowjo version 10 (BD Biosciences). Data was normalized to the average of the WT(s) in each experiment, across 4 different experiments N=6 (all males); females were excluded to due to low numbers (in 1 experiment, n=2 WT, 1 22qDS).

### Immunohistofluorescence of frozen sections

8 μm sagittal brain sections were fixed with ice cold acetone and 70% ethanol, permeabilized with TBS (1X) with 0.025% Tween 20 (Amresco), and nonspecific binding was blocked with 10% normal donkey serum (Sigma) for 4 hours at room temperature. Primary antibodies polyclonal rabbit anti-rat/mouse fibrinogen (1:300; Innovative research), polyclonal rabbit anti-mouse/human claudin-5 (1:300; Life Technologies), polyclonal rabbit anti-mouse/human ZO-1 (1:200; Life Technologies), monoclonal rat anti-mouse ICAM-1 (1:100; Biolegend YN1/1.7.4), monoclonal mouse anti-mouse GFAP (1:2000; Sigma G-A-5), and monoclonal rat anti-mouse IL-6 (1:50; Invitrogen, MP5-20F3) were diluted in 3% normal donkey serum and incubated overnight at 4°C. Sections were washed with TBS (1X), incubated with secondary antibodies diluted 1:300 (Alexa Fluor® 488 AffiniPure F(ab’)_2_ Fragment Donkey Anti-Rabbit IgG, Alexa Fluor® 488 AffiniPure Donkey Anti-Rat IgG, Rhodamine Red™-X (RRX) AffiniPure F(ab’)_2_ Fragment Donkey Anti-Mouse IgG; Jackson Immunoresearch) and Isolectin GS-IB4 From Griffonia simplicifolia, Alexa Fluor® 647 Conjugate (1:300, Life Technologies) in 3% normal donkey serum for 2 hours at room temperature. Nuclei were permeabilized with TBS (1X) with 1:100 Triton X-100 (Amresco) for 10 minutes, and mounted with Gelvatol containing Hoescht nuclear dye (1:1000; BD biosciences). Sections were imaged on Leica widefield microscope (Leica Microsystems) at 10x (GFAP tilescans), 20x (IL-6/GFAP, fibrinogen/IgG) and 40x (claudin-5, ZO-1, ICAM-1). All immunofluorescent analyses were performed blinded and using ImageJ, as previously described (Alvarez et al., 2015). Extravascular fibrinogen and IgG in the anterior cortex was quantified as the average pixel intensity of the leakage area; 12 vessels were analyzed from four 20x z stack maximum projections per animal. Transected vessels stained for claudin-5 and ZO-1 in the anterior cortex were imaged in four 40x z-stack maximum projections per animal. Average pixel intensity was measured at 3 points along each TJ strand and were averaged to a single measurement per TJ; 5 TJ strands from 5 different vessels were analyzed per animal. Vascular ICAM-1 expression was imaged in the anterior cortex in three 40x z-stack maximum projections per animal. Average pixel intensity was obtained at 3 points along each of the four brightest ICAM-1 positive vessels in each animal, and these were averaged to find the average pixel intensity of ICAM-1 per blood vessel. GFAP and IL-6 expression were imaged at 20x in the hippocampus; IL-6 expression was quantified as the average pixel intensity per astrocyte process, with a total of 10 processes analyzed per animal. N=10 (14 females, 6 males from 3 experiments) except assessment of barrier function (fibrinogen/IgG, Figure 1), in which 2 mice noted with poor CNS perfusion at time of brain harvest were excluded.

**Figure 1.**
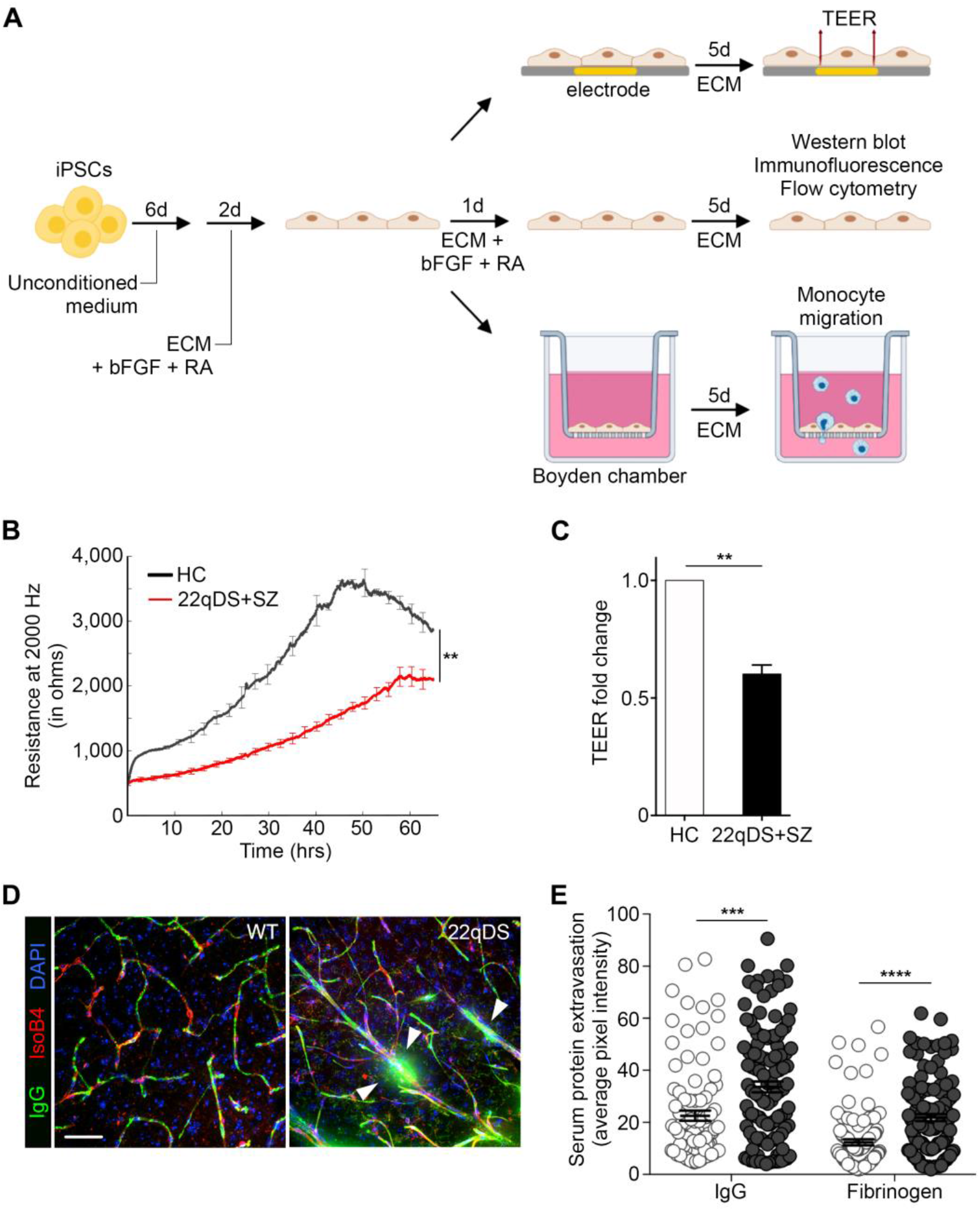
22qDS BBB demonstrates compromised barrier properties. (**A**) iPSCs were differentiated into BBB-like endothelial cells as previously described (UM, unconditioned media; ECM, endothelial cell media; RA, retinoic acid); differentiated cells were grown in ECM + bFGF + RA for 24 hours followed by 5 days culture in ECM prior to experimental analysis. (**B**) Representative TEER curve for one pair of HC and 22qDS+SZ iBBB cells, run in triplicate. (**C**) Quantitative analysis of iBBB TEER data at confluency as a function of fold change across each pair (n = 12, 3 pairs analyzed in 3-5 different experiments). (**D**) Representative immunofluorescent image of IgG (green) in the CNS of WT and 22qDS mice. White arrows indicate extravascular leakage of IgG. Scale bar: 50 μm. (**E**) Quantitative analysis of perivascular extravasation of serum proteins IgG and fibrinogen in the PFC of WT and 22qDS mice (n = 8 WT, 10 22qDS, 12 vessels per animal). Error bars, mean ± SEM. ** *P* ≤ 0.01, *** *P* ≤ 0.001, **** *P* ≤ 0.0001.

### Human post-mortem samples

Using a clinical database of patients with 22qDS, three subjects with full post-mortem neuropathological examinations were identified for inclusion in this study (ages: 1, 2 and 14 months; all patients were female). A pathology database was queried to identify post-mortem samples from age-matched controls. Three controls with congenital heart disease (ages: 1, 1 and 16 months; M/F: 2/1) and two without congenital heart disease (ages: 1 and 12 months; all female) were included. All materials were obtained from pre-existing formalin-fixed, paraffin-embedded tissue of the frontal lobe and were histologically confirmed to contain both cortex and white matter. All materials from human subjects were de-identified in accordance with the Children’s Hospital of Philadelphia Institutional Review Board requirements.

### Immunohistofluorescence of paraffin embedded sections

Paraffin-embedded brains were immunostained as previously published (Alvarez et al., 2002). In brief, sections from post mortem controls (n=3) and 22qDS patients (n=3) were sectioned and mounted on positively charged slides (ThermoFisher Scientific). Sections were deparaffinized by washing in xylene followed by washing in 100%, 95% and 70% ethanol followed by H_2_O. Epitope retrieval was performed in a microwave using citrate buffer pH 6.0 (Dako) for 30 minutes. Tissue sections were permeabilized with wash buffer (Dako) for 5 minutes, and blocked at room temperature for 90 minutes in 10% normal donkey serum (Sigma). Antibodies were diluted in 3% normal donkey serum (monoclonal mouse anti-mouse GFAP 1:2000, Sigma G-A-5; polyclonal rabbit anti-human IL-6 1:50, Proteintech), and incubated overnight at 4°C. Secondary antibody Alexa Fluor® 488 AffiniPure F(ab’)_2_ Fragment Donkey Anti-Rabbit IgG was diluted 1:300 in 3% normal donkey serum, incubated 30 minutes at 37°C, and mounted with Gelvatol containing Hoescht nuclear dye (1:1000; BD Biosciences). Confocal microscopy was performed in an Olympus IX83 as indicated above. IL-6 expression was quantified as average pixel intensity per astrocyte, 15 astrocytes analyzed per patient, using Adobe Photoshop.

### Statistical analysis

Wilcoxon matched-pairs signed rank test was used for comparisons between HC and 22qDS+SZ iBBB experiments. ECIS - TEER statistical analysis was calculated by determining the effect of each genotype on the monolayer of endothelial cells using repeated measures one-way ANOVA. Unpaired student’s t test was used for comparisons between WT and 22qDS mice. All data analysis was performed in GraphPad Prism 7 (GraphPad, San Diego, CA, USA). Values are presented as mean ± SEM. Graphical representations of p values are * *P* ≤ 0.05, ** *P* ≤ 0.01, *** *P* ≤ 0.001, **** *P* ≤ 0.0001.

## RESULTS

### Barrier function is impaired in the 22qDS BBB

As the barrier function of CNS vascular endothelial cells underlies the ability of the BBB to restrict peripheral influences on CNS function (Abbott et al., 2010; Daneman & Prat, 2014), we first assessed barrier properties of the 22qDS BBB. To evaluate the effect of the 22qDS deletion on barrier function, we obtained HiPSCs from 5 22qDS+SZ patients and 5 paired age- and sex-matched HCs. iBBB endothelial cells were derived from these HiPSCs using a differentiation protocol as previously described (Figure 1A) (Hollmann et al., 2017; Lippmann et al., 2012). To interrogate the barrier function of the iBBB monolayers, we assessed their transendothelial electrical resistance (TEER) by repeatedly sampled resistance of the monolayers over the course of 72 hours. We observed a significant decrease in TEER of the 22qDS+SZ iBBB monolayers at confluency compared to paired HC monolayers (Figure 1B), indicating that barrier function of the BBB is compromised in 22qDS (Figure 1C).

To evaluate the 22qDS BBB *in vivo*, we utilized a mouse strain harboring a hemizygous deletion of the 22qDS homologous region on chromosome 16 (Didriksen et al., 2017; Nilsson et al., 2018; Scarborough et al., 2019). These mice mimic much of the biology of 22qDS in humans, including facial deformities and SZ-associated behavioral changes (Didriksen et al., 2017; Nilsson et al., 2018; Scarborough et al., 2019). We assessed BBB integrity by quantifying extravasation of blood proteins into the CNS parenchyma of 22qDS mice and their wild type (WT) littermates. Vasculature was cleared by transcardial perfusion, and immunofluorescent analyses demonstrated a significant increase in extravascular leakage of two serum proteins, immunoglobulin G (IgG) and fibrinogen (Figure 1D-E) in 22qDS. Together, our results support the hypothesis that the BBB is intrinsically compromised in 22qDS.

### Claudin-5 expression is compromised in the 22qDS BBB

As our TEER results indicate impaired junctional integrity underlying compromised barrier function, we next analyzed tight junction protein expression in the 22qDS BBB. Because claudin-5 is the most densely expressed tight junction molecule in the BBB (Morita et al., 1999; Ohtsuki et al., 2007; Ohtsuki et al., 2008) and the gene for claudin-5 is included in the 22q11.2 deletion (Arinami, 2006; Greene et al., 2018), we assessed claudin-5 expression in our iBBB endothelial cell cultures. There was no change in claudin-5 protein level between 22qDS+SZ and HC cultures by western blot (Figure 2 A-B), suggesting that compromised barrier integrity in 22qDS+SZ iBBB monolayers is not solely due to insufficient claudin-5 gene dosage. As the organization of tight junctions is critical for proper barrier function, we assessed claudin-5 localization in the paracellular cleft between endothelial cells, and observed highly disorganized expression in 22qDS+SZ iBBB cells (Figure 2 C-D). This deficit was specific to claudin-5, as we did not observe any changes in the organization of the intracellular tight junction protein zona occludens 1 (ZO-1) (Supplementary Figure 1A).

**Figure 2.**
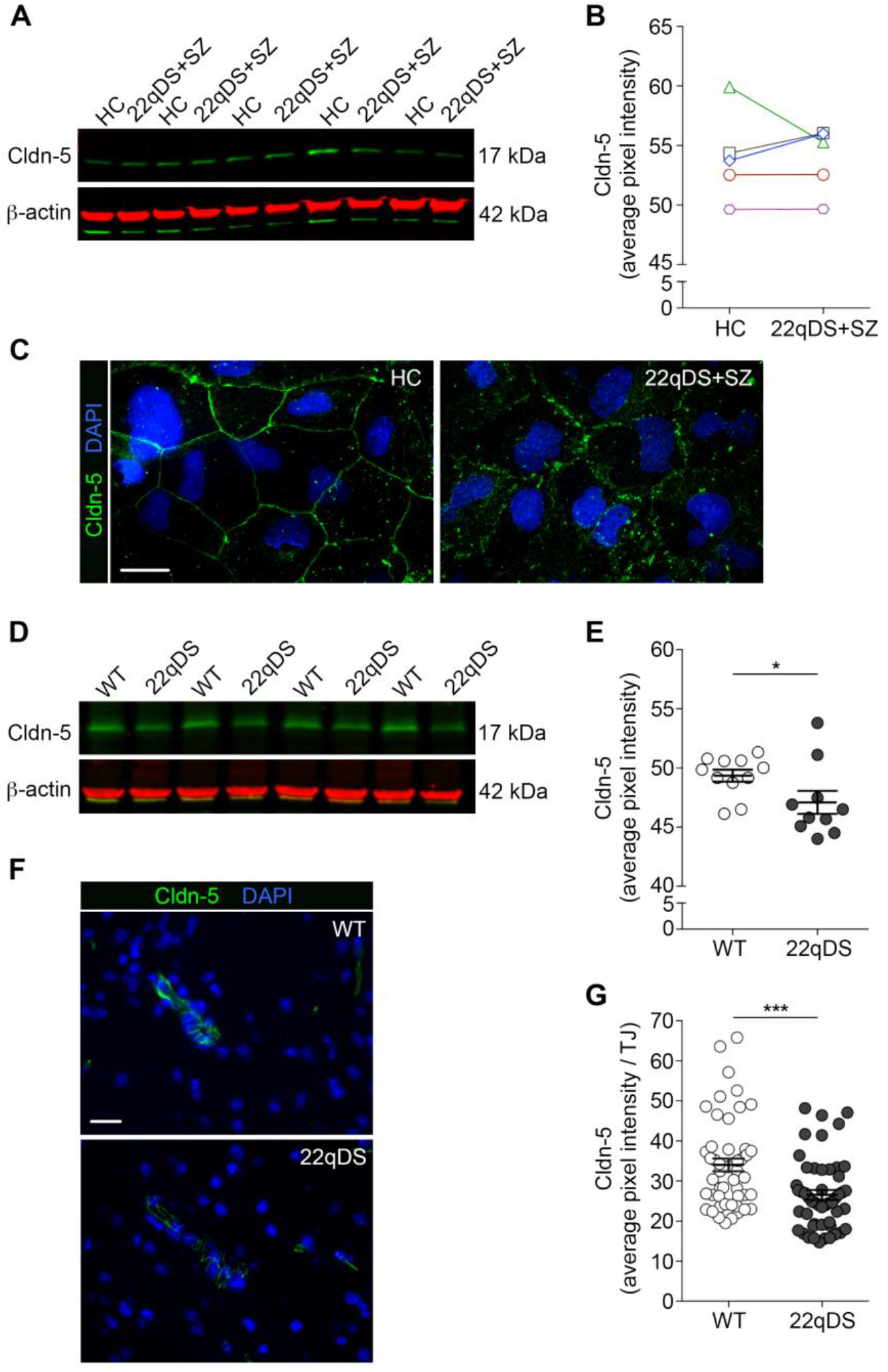
Claudin-5 expression is disrupted in the 22qDS BBB. (**A**) Western blot and (**B**) quantitative analysis of claudin-5 expression levels in HC and 22qDS+SZ iBBB cultures (n = 5, 5 pairs analyzed in one experiment, data represented by colors corresponding to each pair of donors). (**C**) Representative immunofluorescent image of claudin-5 (green) distribution in HC and 22qDS+SZ iBBB cultures. Scale bar: 10μm. (**D**) Representative western blot of claudin-5 in WT and 22qDS mice brain tissue. (**E**) Quantitative analysis of claudin-5 expression level in the PFC of WT and 22qDS mice (n = 11 WT, 10 22qDS). (**F**) Representative 60x immunofluorescent image and (**G**) quantitative analysis of claudin-5 (green) expression in the PFC of WT and 22qDS mice (n = 10, 5 TJs per mouse). Scale bar: 20 μm. Error bars, mean ± SEM. * *P* ≤ 0.05, *** *P* ≤ 0.001.

In contrast to *in vitro* claudin-5 expression, we observed reduced levels of claudin-5 in the brains of 22qDS mice compared to WT by both western blot (Figure 2 D-E) and immunofluorescence (Figure 2 F-G). Again, we did not observe any differences in ZO-1 expression between WT and 22qDS mice, indicating that defects are specific to claudin-5 (Supplementary Figure B-E). Thus, claudin-5 expression is impaired in the 22qDS BBB.

### Immune privilege properties are affected in the 22qDS BBB

Given that the BBB serves as an immunological boundary between the periphery and the CNS (Abbott et al., 2010; Daneman & Prat, 2014; Engelhardt & Coisne, 2011; Muldoon et al., 2013), and because of the renewed focus on the immune system in SZ (Miyaoka et al., 2017; Müller, 2018; Pollak et al., 2018; Severance et al., 2016; Wei & Hemmings, 2005), we assessed the immunoquiescent properties of the 22qDS BBB. We evaluated expression of typical vascular markers (melanoma cell adhesion molecule, MCAM, CD146; thrombomodulin, TM, CD141) (Larochelle et al., 2012; Rochfort & Cummins, 2015; Tran et al., 1996) and those known to be upregulated during inflammation/stress (intercellular cell adhesion molecule 1, ICAM-1, CD54; vascular cell adhesion molecule 1, VCAM-1, CD106) (Alvarez et al., 2015; Daneman & Prat, 2014; Dietrich, 2002; Muldoon et al., 2013). We found no change in vascular marker expression (Supplementary Figure 2A-B) indicating that the iBBB differentiation process was unaffected by the 22q11.2 deletion. However, we found significantly elevated ICAM-1 expression in 22qDS+SZ iBBB endothelial cells compared to HC pairs (Figure 3A-B), suggesting immune activation of the 22qDS BBB endothelium. As expected, we did not observe any change in VCAM-1 expression (Supplementary Figure C), which is no detected on the BBB under homeostatic conditions (J.I Alvarez, personal communication 11/19).

**Figure 3.**
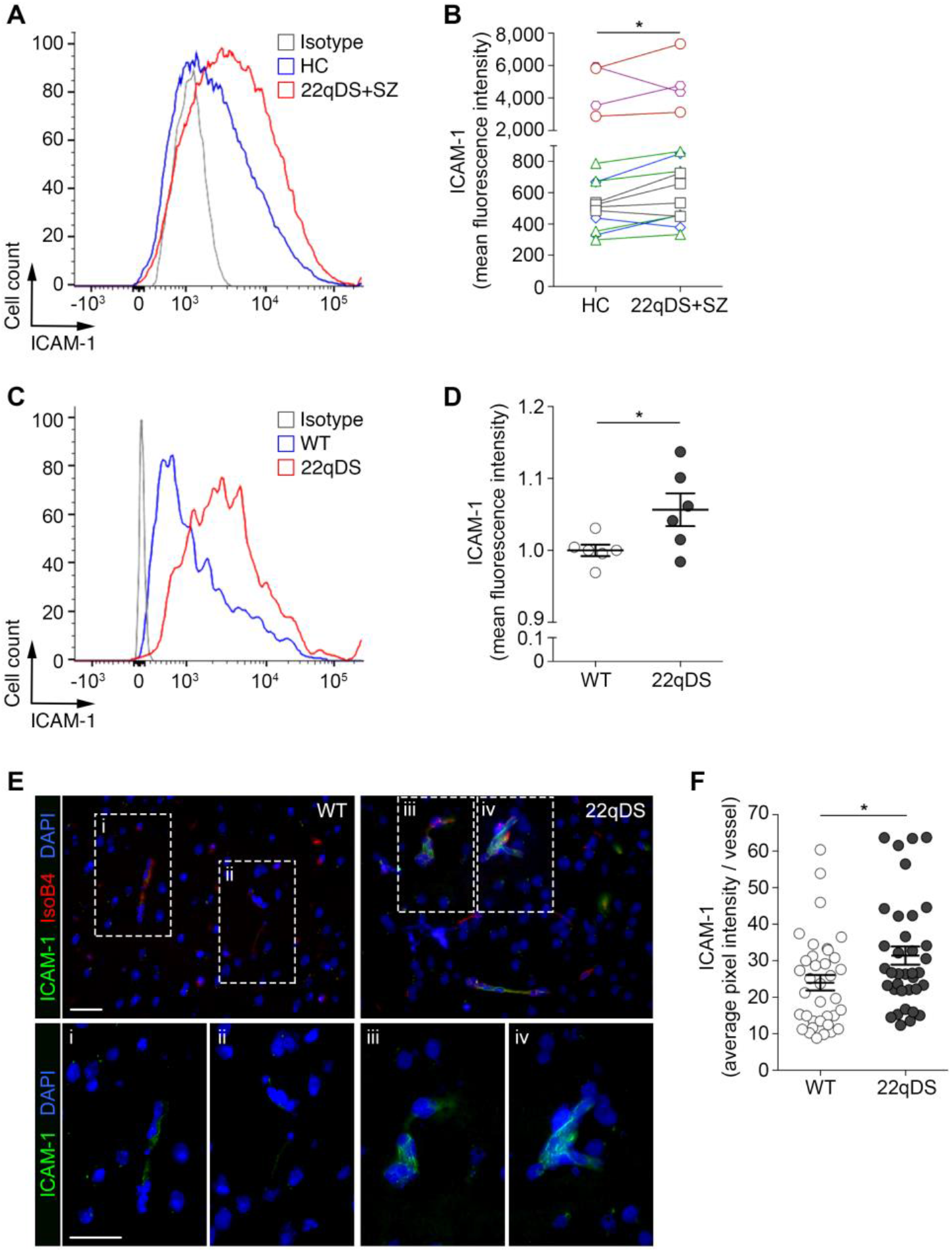
ICAM-1 expression is elevated in the 22qDS BBB. (**A**) Representative histogram plot of ICAM-1 expression on one HC and 22qDS+SZ pair of iBBB endothelial cells. (**B**) Flow cytometry analysis of ICAM-1 expression on HC and 22qDS+SZ iBBB endothelial cells (n = 15, 5 pairs analyzed in 2-4 different experiments, data represented by colors corresponding to each pair of donors). (**C**) Representative histogram plot of ICAM-1 expression on CNS endothelial cells from a WT and a 22qDS mouse. (**D**) Flow cytometry analysis of ICAM-1 expression on CNS endothelial cells from the PFC of WT and 22qDS mice as a function of fold change normalized to WT (n = 6). (**E**) Representative immunofluorescent image at 20x (top) and 60x (bottom) of ICAM-1 (green) expression in vasculature (red) of the PFC in WT and 22qDS mice. (**F**) Quantitative analysis of vascular ICAM-1 expression in the PFC of WT and 22qDS mice (n = 10, 4 vessels per mouse). Scale bar: 20 μm. Error bars, mean ± SEM. * *P* ≤ 0.05.

To assess the immunological properties of the 22qDS BBB *in vivo*, we isolated vascular endothelial cells from the brain of WT and 22qDS mice and determined their cell adhesion molecule expression by flow cytometry. As in our iBBB cultures, we again observed a significant increase in ICAM-1 expression on CNS endothelial cells in 22qDS (Figure 3 C-D). In situ analysis of 22qDS and WT PFC sections further confirmed elevated ICAM-1 expression on the 22qDS BBB, indicating alterations in the inflammatory status of the 22qDS BBB (Figure 3 E-F). Thus, immune privilege properties of the CNS vasculature are impaired in 22qDS.

### 22qDS BBB promotes monocyte migration and activation

To assess the functional consequences of compromised barrier function and immune properties of the 22qDS BBB, we migrated human immune cells across iBBB endothelial cell monolayers (Figure 4A). We found that iBBB monolayers derived from 22qDS+SZ patients were less able to restrict migration of human monocytes (Figure 4B). To determine the effect of transendothelial migration, we assessed expression of monocyte/macrophage markers of activation and migration on transmigrated monocytes (Figure 4C, Supplementary Figure 3). We found that following migration across 22qDS+SZ iBBB monolayer, monocytes significantly decreased thrombomodulin (TM, CD141) expression (Figure 4D-E). As TM is known to be an anti-inflammatory molecule expressed on both immune cells and the luminal endothelium to inhibit immune cell adherence and extravasation (Loghmani & Conway, 2018; van de Ven et al., 2013; Xu et al., 2015), our findings indicate that the 22qDS+SZ BBB has a pro-migratory effect on immune cells.

**Figure 4.**
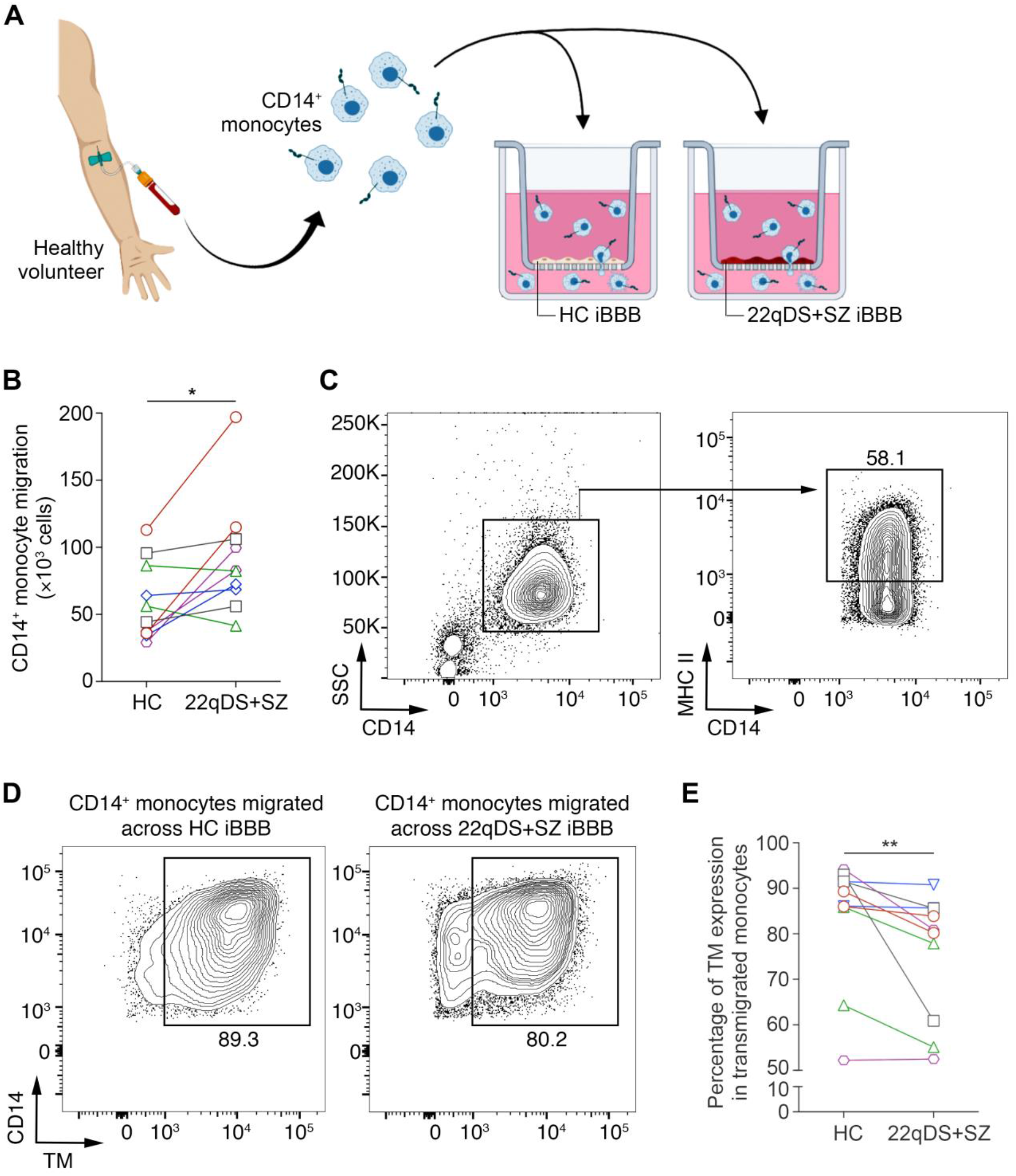
22qDS BBB promotes immune cell transmigration and activation. (**A**) Workflow for isolation and migration of CD14^+^ monocytes. (**B**) Number of monocytes migrating across the iBBB (n = 10, 5 pairs analyzed in 2 different experiments, data represented by colors corresponding to each pair of iBBB donors). (**C**) Gating strategy for analysis of transmigrated CD14^+^ monocytes. (**D**) Representative flow plots of TM expression on transmigrated monocytes. (**E**) Flow cytometry analysis of TM expression on transmigrated monocytes (n = 10, 5 pairs analyzed in 2 different experiments, data represented by colors corresponding to each pair of iBBB donors). * *P* ≤ 0.05, ** *P* ≤ 0.01.

### Perivascular astrocytes are activated in 22qDS

We next aimed to assess the functional consequences of compromised barrier function in 22qDS mice. As astrocytic endfeet ensheath the CNS vasculature (Abbott et al., 2006; Cheslow & Alvarez, 2016), and because we have found that 22qDS mice have elevated perivascular fibrinogen, a molecule that is highly pro-inflammatory in the CNS (Davalos et al., 2012; Schachtrup et al., 2010), we evaluated astrocyte activation by glial fibrillary acidic protein (GFAP) expression. We observed upregulation of GFAP in the brains of 22qDS mice compared to WT, indicating widespread astrocyte activation. Interestingly, this astrocyte activation appeared to be mainly along the CNS vasculature of the meninges (Figure 5A). In order to characterize the activation of astrocytes in 22qDS, we focused our analyses on the pro-inflammatory cytokine IL-6, as this molecule has been repeatedly shown to be elevated in the blood of both idiopathic SZ patients and 22qDS+SZ patients (Khandaker et al., 2014; Mekori-Domachevsky et al., 2017; Potvin et al., 2008; Subbanna et al., 2018a). We found that IL-6 expression was significantly upregulated in astrocytes bordering large CNS vessels of the meninges in 22qDS mice (Figure 5B), suggesting that this activation may not be solely due to intrinsic astrocyte defects, but rather a consequence of compromised barrier function in the 22qDS CNS.

**Figure 5.**
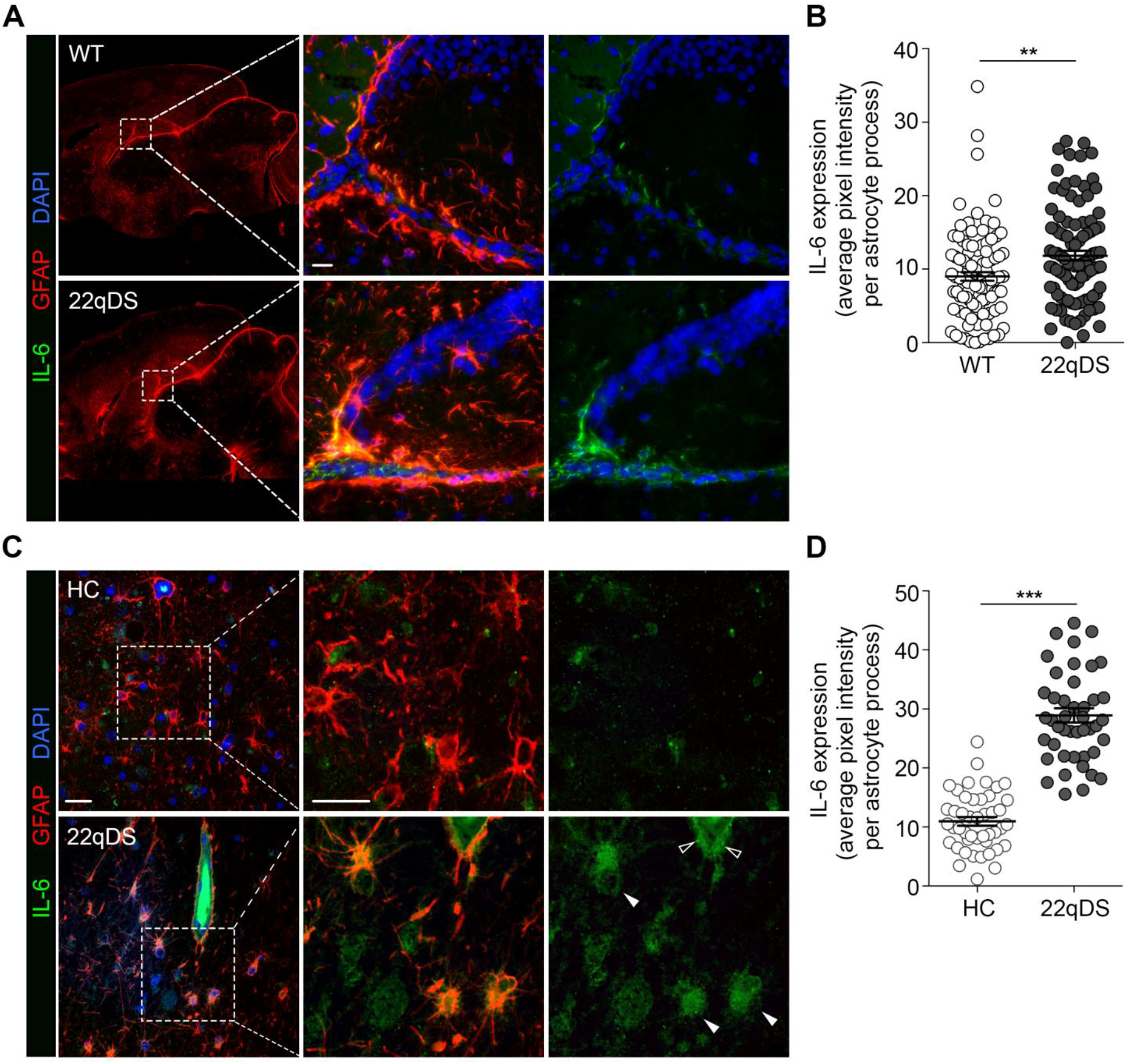
Perivascular astrocytes are activated in 22qDS. (**A**) Representative sagittal sections of WT and 22qDS brains stained for GFAP (red; left, 10x tilescan) and hippocampal meninges stained for GFAP (red) and IL-6 (green) expression (middle, right, 40x). Scale bar: 20μm (**B**) Quantitative analysis of IL-6 expression along hippocampal meninges (n = 10, 10 astrocyte processes per mouse). (**C**) Representative immunofluorescent images of human 22qDS and HC post mortem brain sections stained for GFAP (red) and IL-6 (green) at 20x (left) and 40x (middle, right). White arrows indicate parenchymal IL-6^+^ astrocytes, black arrows indicate perivascular IL-6^+^ astrocytes. Scale bar: 20μm. (**D**) Quantitative analysis of astrocytic IL-6 expression in human 22qDS and HC post mortem brain sections (n = 3, 15 astrocytes per patient). Error bars, mean ± SEM. ** *P* ≤ 0.01, *** *P* ≤ 0.001.

To determine the status of neurovascular astrocytes in human 22qDS patients, we obtained post mortem brain sections from 3 22qDS patients and 3 age-matched HCs. Immunofluorescent staining for GFAP and IL-6 indicated elevated astrocyte activation and IL-6 expression in 22qDS patients compared to HCs (Figure 5C). Together this data further validates that neuroimmune vascular activation in 22qDS involves the astroglial compartment.

## DISCUSSION

For the first time, our results indicate BBB dysfunction in the context of 22qDS, the strongest monogenic risk allele for SZ (Gur et al., 2017). These results are supported by clinical data suggesting that barrier function and immune privilege properties of the BBB are affected in idiopathic SZ patients. Multiple studies have reported that SZ is associated with elevated serum proteins such as immunoglobulins and albumin in the cerebrospinal fluid (CSF), consistent with compromised barrier function (Endres et al., 2015; Severance et al., 2015). Similarly, we have found increased extravasation of serum proteins into the CNS parenchyma in 22qDS mice compared to WT, and this finding of compromised barrier function is further supported by impaired TEER in our 22qDS+SZ iBBB monolayers.

Polymorphisms in the claudin-5 gene have been associated with SZ (Greene et al., 2018; Omidinia et al., 2014; Sun et al., 2004; Wu et al., 2009), including recent work by our lab demonstrating an association between the claudin-5 rs10314 variant and SZ diagnosis in a large cohort female 22qDS patients (Guo et al., 2019). Furthermore, claudin-5 mRNA and protein expression is decreased in the post mortem SZ brain compared to controls (Nishiura et al., 2017). To our surprise, our 22qDS+SZ and HC iBBBs expressed indistinguishable levels of claudin-5 protein, while 22qDS mice expressed significantly less claudin-5 than WT. Rather, altered claudin-5 organization within the paracellular cleft of endothelial cells appears to underlie the compromised barrier function in 22qDS+SZ iBBBs. This contrast between *in vivo* and *in vitro* data suggests that the effects of the 22q11.2 deletion on barrier function are not solely due to claudin-5 gene dosage, but more likely due to an interaction of genes in the deleted region or an exaggerated stress response *in vitro.* Of note are the 6 mitochondrial genes in the 22q11.2 deleted region (Maynard et al., 2008; Napoli et al., 2015), which may further contribute to compromise the BBB given the impaired mitochondrial function reported in other CNS cell populations in 22qDS (Fernandez et al., 2019; Li, Ryan, et al., in press). Furthermore, we have found that ICAM-1 is upregulated in the BBB, implicating differences in inflammatory/stress responses in 22qDS (Alvarez, Cayrol, et al., 2011; Daneman & Prat, 2014; Dietrich, 2002). Alternatively, other factors *in vivo*, including circulating pro-inflammatory molecules and/or the close association with astrocyte endfeet (Abbott et al., 2006; Cheslow & Alvarez, 2016), may alter the effects of the deletion on the BBB *in vivo* compared to *in vitro*.

In addition to compromised barrier function of the BBB, it has also been reported that immunoquiescent properties of the CNS vasculature may be disrupted in SZ patients (Nguyen et al., 2018). Postmortem studies have indicated elevated expression of the pro-inflammatory cell adhesion molecule ICAM-1 in the CNS endothelium of SZ patients (Cai et al., 2018), which may contribute to the increased presence of peripheral immune cells within the CNS parenchyma in post mortem SZ brains (Khandaker et al., 2015). We have also found an increase in ICAM-1 expression in both our 22qDS+SZ iBBB cells and in vascular endothelial cells isolated from the CNS of 22qDS mice. This may contribute to the increase in immune cell migration across the 22qDS+SZ iBBB monolayer. We postulate that the increase in ICAM-1 expression in 22qDS mice may represent an exacerbated immune sensitivity to facilitate leukocyte migration into the CNS upon environmental stress, such as infection or injury.

Using the 22qDS mice, we were able to probe beyond the BBB to interrogate the status of the neurovascular unit. As we had found compromised barrier function *in vivo* and because fibrinogen is known to be a highly pro-inflammatory molecule in the CNS (Davalos et al., 2012; Schachtrup et al., 2010), we hypothesized that this may contribute to astrocyte activation. Our observations build upon previous findings of astrogliosis in the post mortem 22qDS brain (Kiehl et al., 2009) and we report our novel finding that astrocyte activation in 22qDS is associated with significantly increased astrocytic IL-6 expression. Notably, we observed that IL-6 expression was elevated primarily in astrocytes lining the meningeal CNS vasculature in 22qDS mice. Although we cannot attribute our observations of astrogliosis in 22qDS mice specifically to compromised BBB, the pattern of astrocyte activation predominantly along large CNS vessels suggests that impaired barrier function may contribute to astrocyte-mediated neuroinflammation in 22qDS (Muradashvili, Tyagi, & Lominadze, 2017; Schachtrup et al., 2010; Verkhratsky & Nedergaard, 2018). Interestingly, we observed that IL-6 is primarily upregulated by perivascular astrocytes in the hippocampus, a region that has been implicated in clinical SZ patients and behavioral deficits in 22qDS mice (Kalmady et al., 2017; Lieberman et al., 2018; Sauras et al., 2017; S. Anderson, personal communication 11-2019). These results suggest potential mechanisms by which compromised BBB may contribute to the neuronal dysfunction that has previously been reported in 22qDS mice, and future studies could address a causal relationship between BBB impairment and altered neuronal signaling.

Although our approach of utilizing HiPSC lines derived from 5 different 22qDS+SZ human donors introduces more variability into our data than typical *in vitro* studies, our consistent results despite our use of genetically different cell lines further reinforces our conclusions that barrier function and immune properties of the BBB are disrupted in 22qDS+SZ. It is important to note that in both our human 22qDS+SZ iBBB cultures and our human 22qDS+SZ brain sections, we cannot determine the role of SZ diagnosis in our results. It is unclear whether all 22qDS patients present with compromised barrier function, impaired claudin-5 structure, and elevated ICAM-1 expression, which together may contribute to the increased risk for neuropsychiatric conditions in this population, or whether these results are only evident in 22qDS patients that develop SZ. As our murine data indicate BBB dysfunction in naïve 22qDS mice, we hypothesize that our results are conferred by the deletion alone, regardless of SZ diagnosis, and therefore may contribute to the increased susceptibility to SZ and other neuropsychiatric disorders in 22qDS (Fiksinski et al., 2018; Fiorentino et al., 2016; Gandal et al., 2018; Kealy et al., 2018; Schneider et al., 2014), but future studies including a 22qDS without SZ group will be important to address these questions.

This is the first study to assess the status of the BBB in the context of 22qDS, a population known to have a 25% increased risk of developing SZ (Arinami, 2006; Karayiorgou & Gogos, 2004; Van et al., 2017). Our results support the hypothesis that neuroinflammation and compromised BBB may play a role in the pathogenesis of neuropsychiatric disorders. Fundamental to this hypothesis is the clinical data indicating peripheral inflammation in a subpopulation of SZ patients, including elevated peripheral cytokines (most notably IL-6) and immune cell activation (especially of Th17 cells) (Debnath & Berk, 2014; Potvin et al., 2008; Subbanna et al., 2018b). Interestingly, 22qDS patients present with a similar inflammatory profile, as they also have elevated peripheral IL-6 and increased percentages of circulating Th17 cells (Mekori-Domachevsky et al., 2015; O’Rourke & Murphy, 2019; Vergaelen et al., 2018). Furthermore, a key component to the association between this reported immune activation and SZ is that BBB disruption facilitates CNS inflammation and neuronal dysfunction. Our results suggest that 22qDS may represent a uniquely appropriate condition to assess the role of peripheral inflammation and BBB dysfunction in neuropsychiatric disorders like SZ.

## ACKNOWLEDGEMENTS

The National Institute of Health (NIH) of the United States has supported this work by the following grants NIMH 5R01MH110185-03 (SAA) and NINDS 5K01NS097519-03 (JIA).

We want to thank Drs. Raquel Gur and Tim O’Brian and the Neurobehavioral Core at the University of Pennsylvania for maintenance of the 22qDS mouse colony. We also want to thank Dr. Douglas Coulter and his laboratory for sharing reagents to carry out this study and the laboratory of Dr. Herbert M. Lachman, MD (Albert Eistein College of Medicine) for providing the HiPSC lines used in this study. We appreciate the advice obtained from Gordon Ruthel at the Penn Vet Imaging Core at the University of Pennsylvania for advice on scoping and image analysis.

---

**Supplementary Figure 1.**
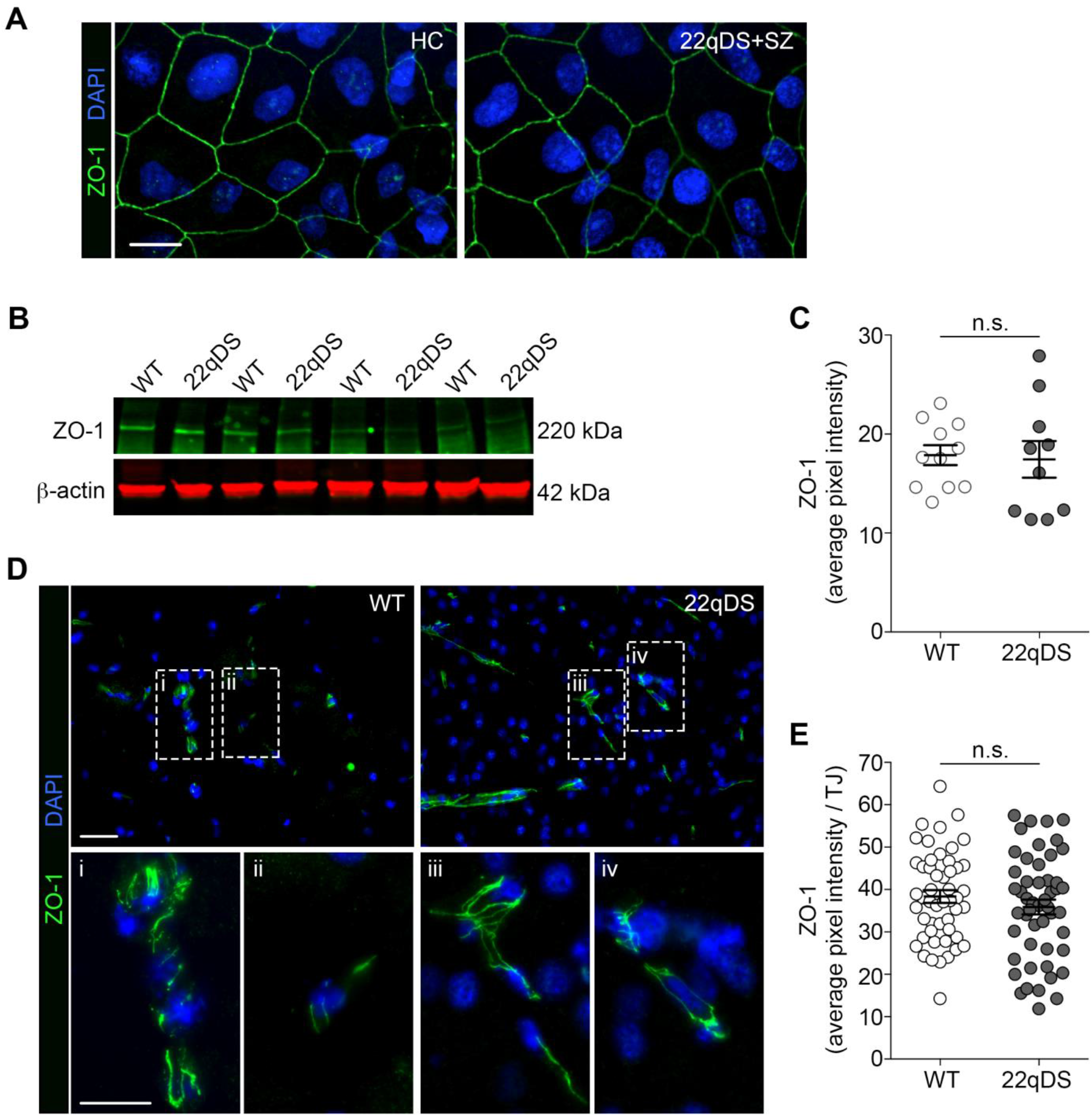
ZO-1 expression is unchanged in the 22qDS BBB. (**A**) Representative immunofluorescent image of ZO-1 (green) distribution in HC and 22qDS+SZ iBBB cultures. Scale bar: 10μm. (**B**) Representative western blot of ZO-1 in WT and 22qDS mice brain tissue. (**C**) Quantitative analysis of ZO-1 expression level in the PFC of WT and 22qDS mice (n = 11 WT, 10 22qDS; *P* = 0.83). (**D**) Representative 60x immunofluorescent image and (**E**) quantitative analysis of ZO-1 (green) expression in the PFC of WT and 22qDS mice (n = 10, 5 TJs per mouse; *P* = 0.28). Scale bar: 20 μm. Error bars, mean ± SEM.

**Supplementary Figure 2.**
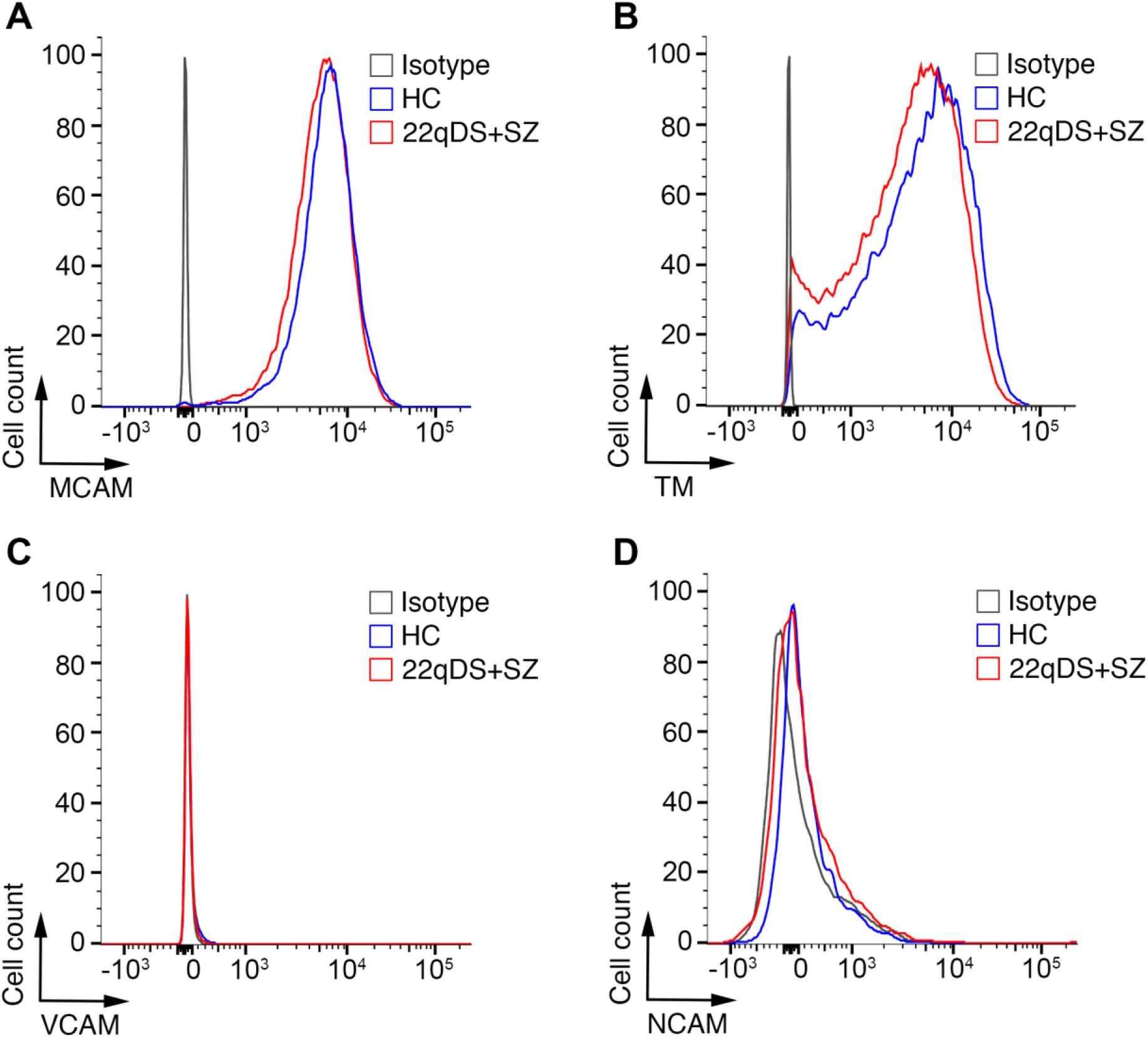
22qDS+SZ iBBBs expression of CNS vasculature markers and cell adhesion molecules are unchanged from HC. (**A**) MCAM (CD146) and (**B**) TM (CD141) are expressed by our iBBB cultures, indicated vascular phenotype; expression is not different between 22qDS+SZ and HC pairs. (**C**) VCAM (CD106) is not expressed by 22qDS+SZ and HC pairs, as VCAM is rarely expressed on the BBB during homeostasis. (**D**) Neural cell adhesion molecule (NCAM; CD56) is not expressed by 22qDS+SZ and HC pairs, indicating non-neuronal, non-glial phenotype as expected.

**Supplementary Figure 3.**
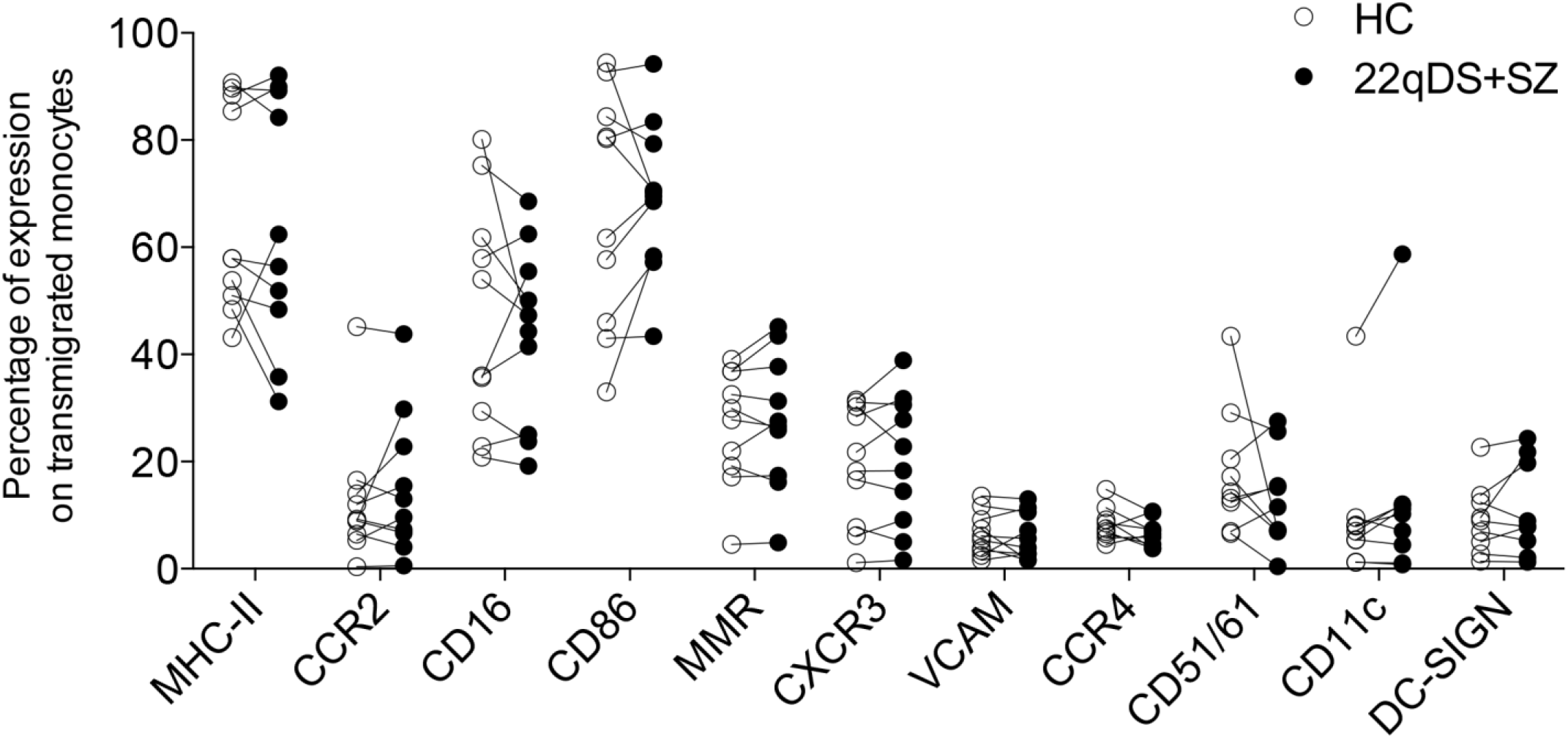
Additional monocyte/macrophage surface markers are unchanged by migration across 22qDS+SZ iBBB. Flow cytometry analysis of activation and migration markers expressed on transmigrated monocytes (n=10, 5 pairs analyzed in 2 different experiments, data represented as paired iBBB healthy control (HC) donors). MHC-II, *P* = 0.43; CCR2, *P* = 0.50; CD16, *P* = 0.36; CD86, *P* = 0.43; MMR (CD206), *P* = 0.60; CXCR3 (CD183), *P* = 0.62; VCAM (CD106), *P* = 0.55; CCR4 (CD194), *P* = 0.16; CD51/61, *P* = 0.30; CD11c, *P* = 0.20; DC-SIGN (CD209), *P* = 0.65.

